# Sex-Specific Protection Against Diet-Induced Non-Alcoholic Fatty Liver Disease in TRPV1 Null Mice

**DOI:** 10.1101/403881

**Authors:** Patrick J. Connell, Ian N. Bratz, Spencer R. Andrei, Luke Eusebio, Daniel J. DelloStritto, Joseph N. Fahmy, Jessica M. Ferrell, Preeti Pathak, John Y.L. Chiang, Derek S. Damron

**Affiliations:** Department of Integrated Medical Sciences, Northeast Ohio Medical University, Rootstown, OH 44272; Department of Medicine, Vanderbilt University Medical Center, Nashville, TN, 37217; Department of Biological Sciences, Kent State University, Kent, OH, 44242

**Author notes:** Underlined name indicates equal contribution (Patrick J. Connell, Ian N. Bratz, Spencer R. Andrei). **Corresponding Author:** Derek S. Damron, Ph.D., Dept. of Biological Sciences/Cunningham Hall, Kent State University, Kent, OH 44240.

**Keywords:** Obesity, TRPV1, Diabetes, NAFLD, Liver

## Abstract

**Objectives:** TRPV1 channels have been linked to the development and progression of diabetes at multiple levels, including control of appetite and weight, regulation of pancreatic function, thermogenesis, metabolism and energy homeostasis. Despite this, little information is known regarding its role in liver homeostasis and nonalcoholic fatty liver disease (NAFLD).

**Methods and Results:** To better understand the role of TRPV1 in liver metabolism, we explored the effects of a high fat/sugar diet (Western, 24-week regimen) in male and female wild type (WT) and TRPV1-null (V1KO) mice. Our data reveal that loss of the TRPV1 gene makes mice susceptible to diet-induced obesity and induces NAFLD. V1KO mice displayed gross phenotypic and gross morphological changes including insulin resistance, glucose intolerance, increased body mass and central adiposity on a western diet compared to WT counterparts. Western fed V1KO mice exhibited gross changes in liver morphology and size compared to western fed WT mice, which were supported with histological H&E and Oil Red O staining. Accompanying the liver changes, Western fed V1KO mice exhibited altered lipid profiles as demonstrated by elevated hepatic triglyceride, cholesterol and free fatty acid levels compared to western fed WT mice. Interestingly, female V1KO mice fed a western diet displayed significant protection against diet-induced obesity and the progression of NAFLD compared to their male counterparts. Taken together, these data suggest that loss of TRPV1 promotes fat accumulation, NAFLD development and changes in liver lipid profiles in male mice, the extent to which is less severe in female V1KO mice.

**Conclusion:** In conclusion, TRPV1 may be a protective therapeutic target for the prevention of NAFLD development in diet-induced obesity.

## 1. Introductory Statement

Diet-induced obesity is an epidemic that predisposes individuals to cardiometabolic complications, including type 2 diabetes mellitus (T2DM) and nonalcoholic fatty liver disease (NAFLD), which are conditions directly related to inappropriate ectopic lipid deposition, fatty deformation, and altered hepatic lipid metabolism. NAFLD refers to hepatic steatosis, or the accumulation of lipid in the liver, not related to alcohol consumption (1). The prevalence of NAFLD has been reported to be as high as 30% in western countries (2). NAFLD prevalence is directly correlated with obesity and insulin resistance. As such, current trends in obesity and diabetes epidemiology predict an increased incidence of NAFLD worldwide. NAFLD includes a broad spectrum of diseases from simple hepatic steatosis to inflammatory steatohepatitis with increasing levels of fibrosis and ultimately cirrhosis (3). The pathogenesis of NAFLD and its progression to fibrosis and chronic liver disease are still unknown. A popular “two-hit” hypothesis has been proposed in which the first hit is an initial metabolic alteration, such as insulin resistance, hyperglycemia, and the accumulation of triglyceride in hepatocytes, leading to hepatic steatosis. The second hit induces the progression to more severe injury including steatohepatitis, inflammation, fibrosis, and cirrhosis (4). These excesses lead to adipogenesis and adipocyte differentiation in both subcutaneous and visceral areas and tissue, which have been shown to disturb regulation of lipid protein signalers such as cytokines, chemokines, and adipocytokines, resulting in impaired lipid signaling, lipid metabolism and systemic inflammation. Altered lipid metabolism in particular occurs through factors that are linked to adipose tissue, such as free fatty acids, leading to a hindered metabolic state via increased hepatic insulin resistance and fat accumulation in the liver.

Currently, the principal therapeutic modalities for NAFLD are lifestyle interventions (5), while pharmacologic treatments used for improving hepatic inflammation, fibrosis, and clearing steatohepatitis offer limited results (6). Recent evidence indicates that activation of <u>T</u>ransient <u>R</u>eceptor <u>P</u>otential <u>V</u>anilloid <u>1</u> (TRPV1) cation channels by its agonist, capsaicin, is beneficial for the management of obesity, diabetes and related diseases. TRPV1, a polymodal cation channel, has been found to be expressed ubiquitously. Human and animal studies support a requisite role for TRPV1 and capsaicin to regulate energy expenditure and metabolism, reduce adipose tissue weight and increase lipid oxidation (7). Similarly, TRPV1 has also been shown to regulate adipocyte function through the utilization of calcitonin gene-related peptide (CGRP)-dependent mechanism, to thus prevent preadipocyte differentiation (8, 9). As such, capsaicin and TRPV1 could potentially be a useful strategy to combat diet-induced obesity and cardiometabolic diseases such as NAFLD.

A role for TRPV1 in the physiology and pathophysiology of liver function is beginning to be elucidated, but there is much more that remains to be understood. For instance, TRPV1 activation has been shown to modulate the inflammatory responses and fatty acid oxidation in the liver (10). Li et al., 2012 demonstrated that TRPV1 activation leads to increased hepatic uncoupling protein 2 (UCP2), to elicit a reduction in fatty acid content in the hepatocyte mitochondrial matrix and thus an overall decrease in hepatic fatty deposition (11). In this way, it is evident that TRPV1 improves hepatic health and prevents NAFLD, yet the specific mechanistic roles of TRPV1 in the liver are still unknown. In the current study, we present experimental evidence that loss of TRPV1 channels contributes to diet-induced obesity and the development of T2DM and NAFLD in mice fed a high fat/sugar diet for 24 weeks.

## 2. Experimental Procedures

### 2.1. Animal Models

All procedures were conducted with the approval of the Institutional Animal Care and Use Committee of the Northeast Ohio Medical University (NEOMED) and in accordance with National Institutes of Health Guidelines for the Care and Use of Laboratory Animals. Breeding pairs of mice were originally purchased from Jackson Labs (Bar Harbor, ME) after which mice were bred in the NEOMED animal facility. Experiments were performed in 5 and 29-week-old male and female C57BL/6J (WT) mice and TRPV1^−/−^ (V1KO) mice. Unless otherwise indicated, mice were housed individually in a temperature-controlled room (23°C) in virus-free facilities on a 12-h light/dark cycle (7:00 a.m. on/7:00 p.m. off) and were fed a standard rodent chow (No. 5001, Test Diet) and water *ad libitum* for the first 5 weeks. At 5 weeks of age, both male and female WT and V1KO mice were randomly assigned into respective groups and fed either a standard chow (Chow) or western diet (Western, Harlan TD 88137; 42% fat, 42.7% carbohydrate, 15.2% protein, 0.2% cholesterol) for 24 weeks; the four groups (n = 10/group) consisted of; WT Chow (CWT), WT western diet (WWT), V1KO Chow (CV1KO) and V1KO Western (WV1KO)). On the last day of the study, blood was collected from the orbital plexus and mice were then subjected to cervical dislocation. Organs were harvested, weighed, and processed for histological analysis, quantitative RT-PCR and Western blotting respectively. Images of livers were taken for qualitative analysis. Tissues not used for histology were snap frozen in liquid nitrogen and stored at −80°C until further use.

### 2.2. Metabolic Analyses

Metabolic activity and respiratory quotient were determined in both 5 week-old and 29 week-old WT and V1KO mice (maintained on chow or Western diet) using a Comprehensive Lab Animal Monitoring System (CLAMS) (Columbus Instruments; Columbus, OH); a series of live-in cages for automated, non-invasive and simultaneous monitoring of food and water consumption; horizontal and vertical activities and metabolic performance. O_2_ consumption (VO_2_), CO_2_ production (VCO_2_), respiratory exchange ratio (RER [VCO_2_/VO_2_], an estimate of fuel usage) and energy (heat) production were calculated and recorded electronically over 24 hours for each mouse (following a 48-hour acclimation period).

### 2.3. Metabolic Assessment

Weekly diet consumption and body weight were recorded in all mice. In conjunction, Echo Magnetic Resonance Imaging (EchoMRI 3-in-1 Body Composition Analyzer; Houston, TX) was performed to assess measurements of fat and lean tissue mass in live mice following an overnight fast. Fasting plasma glucose samples were collected via tail vein using a Contour Next EZ glucometer (Bayer; Mishawaka, IN).

### 2.4. Glucose Tolerance

Glucose tolerance tests (GTT) were performed in separate cohorts of both 5 week-old and 29 week-old (maintained on chow or Western diet) male and female WT and V1KO mice by intraperitoneal injection of D-glucose (1 g/kg of body weight). Blood samples were taken at 0, 15, 30, 60, 120 minutes from the tail vein. Blood samples were collected at 0, 15, 30, 60, 120 minutes via tail vein. Plasma glucose was measured using a Contour Next EZ glucometer.

### 2.5. Hepatic Lipid Extraction and Histology

Lipid extraction from liver was performed as described (12). Extracted TG and cholesterol contents were normalized to wet liver weight. Commercially available assays were utilized to measure the concentration of triglycerides (Infinity, ThermoFisher), cholesterol (Infinity) and free fatty acids (Wako Diagnostics, Richmond, VA) in liver tissue and serum. 100 mg liver samples were homogenized and lipids were isolated using a 7:10:0.1 ratio of chloroform:isopropanol:NP-40. Serum was prepared after centrifugation of whole blood at 12,000 *g* for 10 min. For histology, livers collected from mice were embedded in Tissue-Tek OCT compound. After sectioning, samples were stained with hematoxylin and eosin (H&E) or Oil Red O for microscopic observation. The sections were visualized under an Olympus upright microscope fitted with a SPOT-imaging camera. The images were then enhanced and color corrected using Autodesk Pixlr Editor.

## 3. Results

### 3.1. V1KO mice are susceptible to diet-induced phenotypic changes

At 5 weeks of age, mice exhibited no differences in body weights or lean mass to body weight ratios (LM/BW) between groups. Wild type and V1KO mice exhibited normal weight gain and growth on the control diet. Interestingly, both male and female V1KO mice on the western diet exhibited a significant increase in body weight over the 24-week period, in contrast to male and female WT mice fed the western diet (Fig. 1A and 1B). Male V1KO mice displayed a notable increase in central white fat normalized to BW compared to their WT counterparts regardless of diet, yet this effect was more pronounced following diet-induced obesity (Fig. 1C). In contrast, female V1KO mice only exhibited an increase in central adiposity following the western diet regimen (Fig. 1D). Using Echo MRI, lean mass to body weight compositions were collected at 5 and 29 weeks of age. No differences in lean mass to body weight ratio were observed between wild-type and knockout mice at baseline. However, after 24 weeks of diet-induced obesity, both WT and knockout mice exhibited a significant decrease in this ratio, which was further exacerbated in V1KO animals (Fig. 1E). This decrease in LM/BW correlated with an increase in overall fat mass (Fig. 1F). The differences in these observations between sexes indicate that female V1KO mice are comparable to wild-type mice on a normal diet, while male V1KO mice are pre-dispositioned to dysfunction without Western diet.

**Fig. 1:**
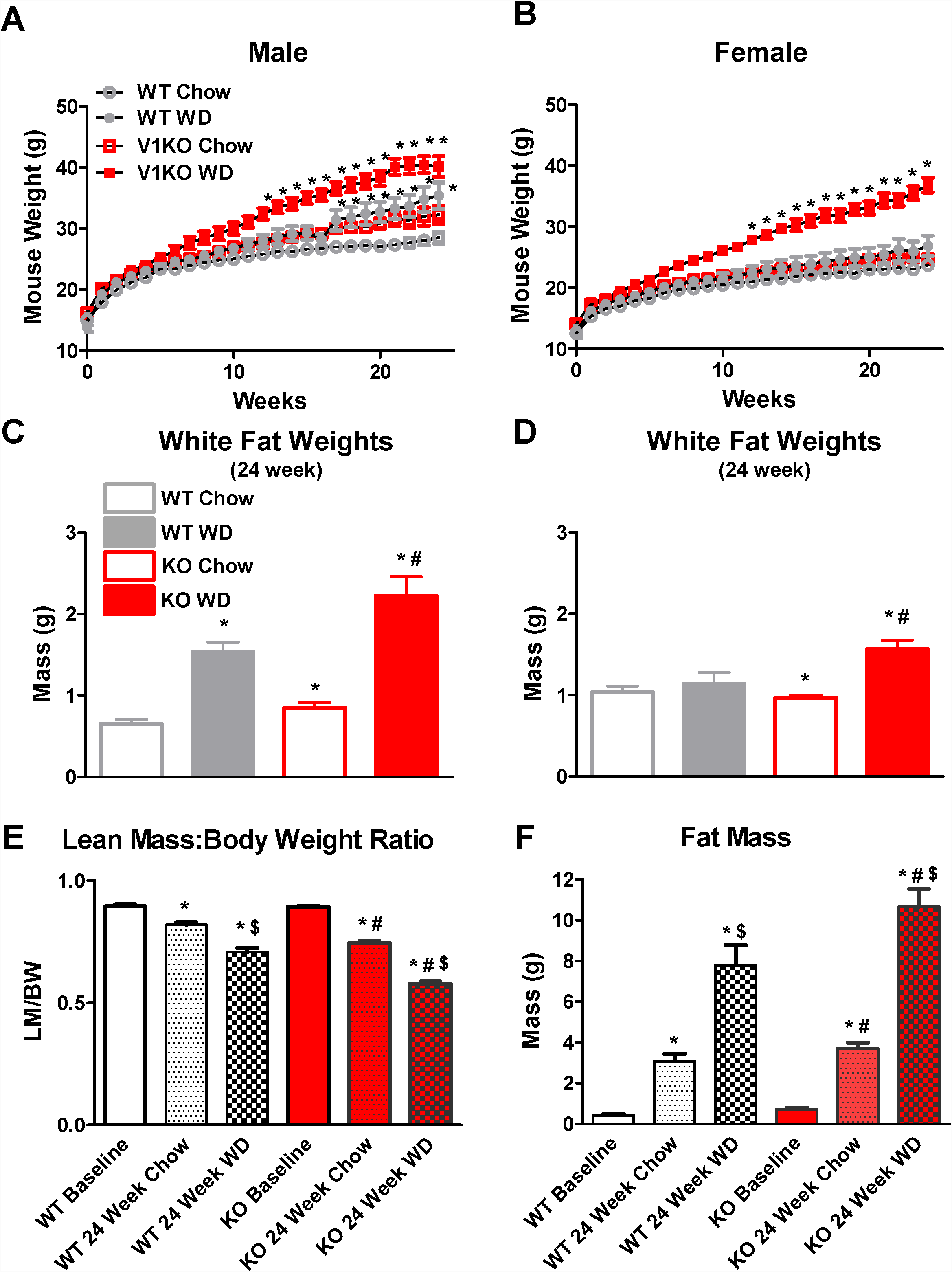
V1KO mice are susceptible to diet-induce obesity. Growth curve of male (A) and female (B) WT and TRPV1 knockout (KO or V1KO) mice fed standard rodent chow (4% fat; ≤0.04% cholesterol) or western diet (WD; high-fat, high-sugar diet containing 42% fat, 42.7% carbohydrate, 15.2% protein) for 24 weeks. Central adiposity of male (C) or female (D) WT and V1KO mice fed standard rodent chow or WD for 24 weeks. (E) Combined male and female lean mass to body weight ratios (LM/BW) in WT and V1KO mice at 5 and following 24 weeks of WD. (F) Combined male and female fat mass in WT and V1KO mice at 5 and following 24 weeks of WD. All values are expressed as mean ± SEM. **\****P* < 0.05 vs. WT chow analyzed by two-way ANOVA. *n* = 10 mice for each treatment group.

### 3.2. V1KO mice exhibit severe glucose intolerance after Western diet regimen

To examine if diet-induced obesity is associated with altered glucose homeostasis, differences in fasted glucose levels and glucose handling (as measured by GTT) were measured in all groups at 5 and 29 weeks of age. At 5 weeks of age, fasting glucose levels showed no difference between groups. Following completion of the 24-week diet regiment, both male and female western diet fed V1KO mice exhibited a significant increase in fasted glucose compared to their WT counterparts (Fig. 2A and 2B). To further assess glucose handling, GTTs were performed in 29 week mice feed chow or western diet. Regardless of diet, both male and female V1KO mice exhibited glucose intolerance, while only the western diet fed WT mice exhibited this intolerance (Fig. 2C and 2D).

**Fig. 2:**
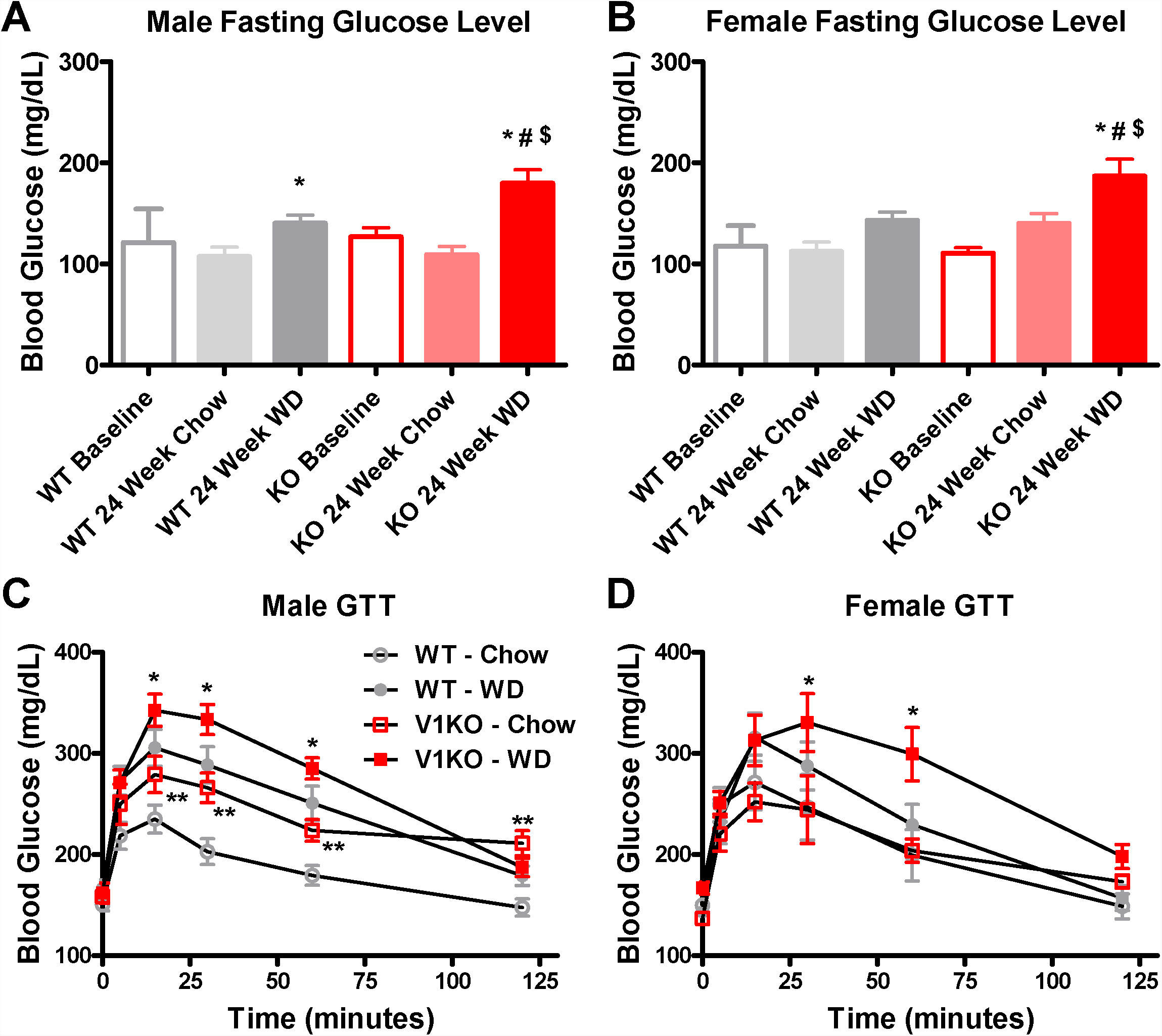
V1KO mice exhibite altere metabolic indices following 24 week W regimen. Fasting glucose levels of male (A) and female (B) WT and V1KO mice fed standard rodent chow or WD for 24 weeks. Glucose tolerance test (GTT) in male (C) and female (D) WT and V1KO mice fed standard rodent chow or WD for 24 weeks. All values are expressed as mean ± SEM. **\****P* < 0.05 vs. control analyzed by two-way ANOVA. #*P* < 0.05 vs. group analyzed by two-way ANOVA. *n* = 10 mice for each treatment group.

Using CLAMS metabolic assessment, RER was determined in mice at 5 weeks and following 24 weeks of diet. At 5 weeks, RER was significantly increased in knockout male mice compared to male WT counterparts (Fig. 3A). Following 24 weeks of high fat diet, RER was significantly decreased in both male groups compared to their counterparts, which were fed standard chow for 24 weeks, yet no significant difference was seen between male groups fed Western diet for 24 weeks (Fig. 3B). Parallel experiments were conducted in WT and KO female mice. No statistical differences in RER were observed between WT or KO mice at either the 5 or 24-week post-diet time points (Fig. 3C and Fig. 3D, respectively).

**Fig. 3:**
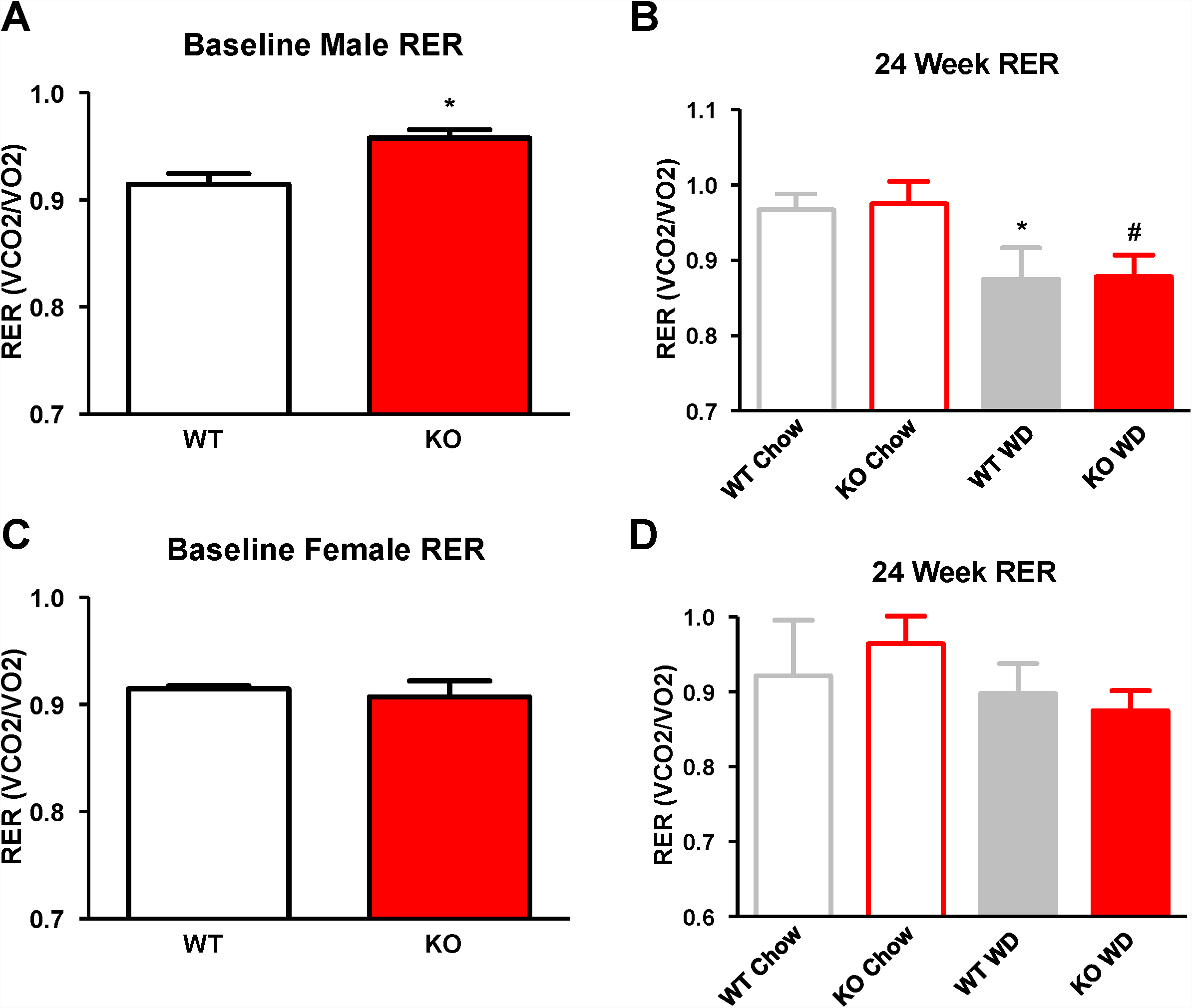
CLAMS metabolic assessment. CLAMS assessment of respiratory exchange ratio (RER) values at 5 weeks of age in male (A) and female (C) WT and KO mice. RER values in male (B) and female (D) mice following 24 weeks period of mice fed either standard rodent chow or WD. All values are expressed as mean ± SEM. **\****P* < 0.05 vs. WT chow. #*P* < 0.05 vs. KO chow. *n* = 6 each group.

### 3.3. Hepatotoxicity and lipid accumulation in TRPV1-deficient Mice

In agreement with the gross phenotypic changes related to diet-induced obesity, alterations in liver morphology revealed the deleterious effects of long-term western feeding in both groups. Diet-induced obesity resulted in a marked increase in liver weight in both wild-type and V1KO mice, which was more prominent in male V1KO mice (Fig. 4A-E). Paralleling this effect, livers from V1KO mice were paler in color with a considerable increase in size and liver to body weight ratios (Fig. 4A-D). Chow-fed V1KO male mice also had larger livers than chow-fed matched WT mice. In female mice of both groups, liver morphology was protected against diet-induced obesity in comparison to male counterparts (Fig 4E-H). To further assess the effects of diet-induced obesity on liver hepatotoxicity and lipid accumulation, histology was assessed in liver sections using H&E and Oil Red O staining. Oil Red O staining revealed a consistent increase in neutral lipid content of the V1KO livers on both diets, including the control chow. As expected, staining was increased in both wild type and V1KO livers on the chow and western diets, and revealed dramatic neutral lipid accumulation in livers obtained from the V1KO Western-fed diet animals (Fig. 5A-D).

**Fig. 4:**
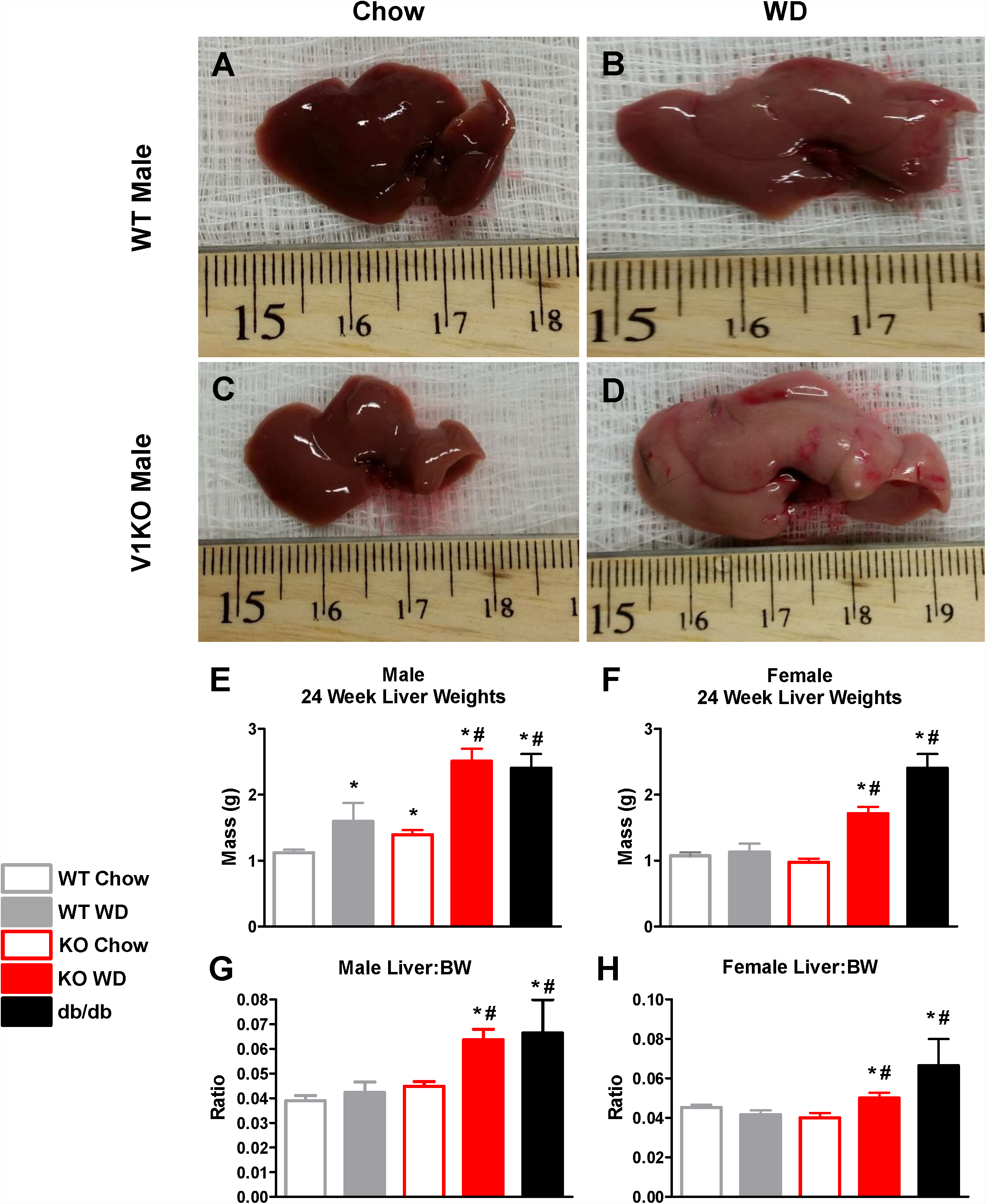
Hepatotoxicity, liver damage an lipi accumulation in TRPV1-deficient mice following 24 week diet regimen. Gross hepatic morphological changes in wild-type male mice fed either chow (A) or western (B) diets and in V1KO male mice fed chow (C) or western (D) diets. Male (E) and female (F) liver weights following 24 weeks of diet. Male (G) and female (H) liver mass to body weight ratios following 24 weeks of diet in WT and V1KO mice. All values are expressed as mean ± S.E. **\****P* < 0.05 vs. WT chow. #*P* < 0.05 vs. KO chow. *n* = 10 mice for each treatment group except db/db (*n* = 5).

**Fig. 5:**
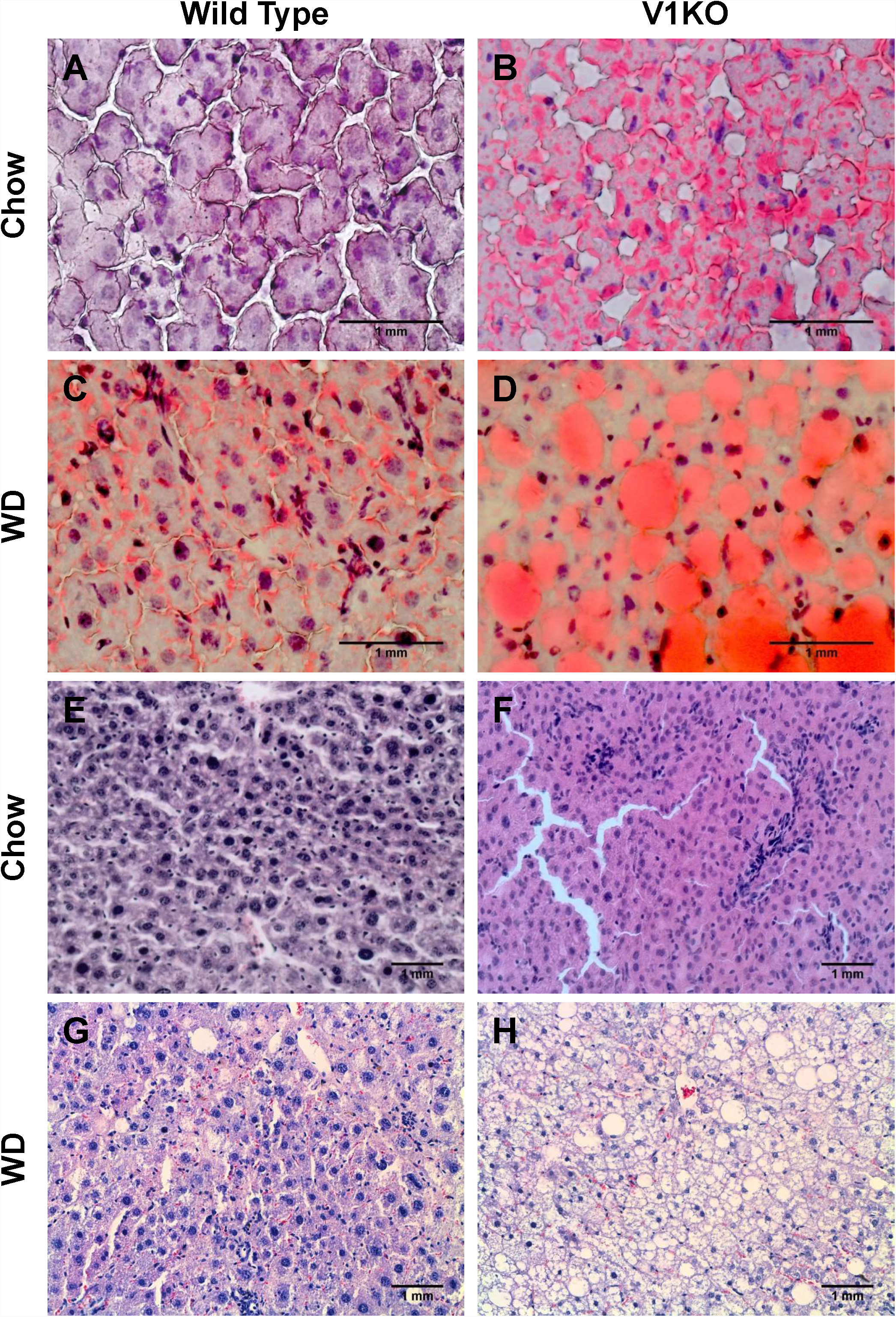
Hepatic fat accumulation in V1KO mice challenge with diet-induce obesity. Male WT and V1KO were placed on the chow or WD regimen. After the dietary intervention, mice were euthanized for collection of livers. *(A-D)* Representative Oil Red O-stained liver sections (×40). Lipid-containing vacuoles stained *red*. Lipid accumulation is significantly higher in the V1KO animals on WD. (E-F) Representative hematoxylin and eosin–stained liver sections (x20) from WT and V1KO mice were examined following 24 week diet regimen. Enlarged lipid depots in degenerating hepatocytes are more evident in WD fed V1KO mice than WT on WD. Features of liver pathology, including inflammation are much more severe in V1KO mice on the WD. Degeneration of individual hepatocytes, loosened liver structure, increased cell size, enlarged sinusoidal spaces, and accumulation of lipid droplets are seen in V1KO mice on the WD, but to a lesser extent in WT mice on WD.

Obvious differences were observed between WT and V1K mice on the Western diet using H&E staining. Hepatocyte damage, inflammation or disruption of overall architecture and increased lipid storage were clearly observed (Fig. 5E-H). Further evidence of hepatocellular degeneration in livers of wild type and V1KO mice on the western diet was also observed. In striking contrast, the livers of WT mice were mildly affected by western diet, with degeneration of individual hepatocytes along with increases in cell size and intercellular sinusoidal spaces (Fig. 5E-H). These effects were much more prominent in V1KO mice (Fig. 5E-H). Thus, histological analysis confirms the increased toxicity and lipid accumulation as a result of high fat diet consumption in the V1KO mice.

### 3.4. Altered Lipid Homeostasis in TRPV1-Deficient Mice

To further confirm morphological and histological effects on lipid accumulation in the liver, hepatic lipid profiles (cholesterol, triglyceride, and free fatty acid) were examined in each group. The hepatic cholesterol, triglyceride and free fatty acid levels of V1KO mice were all significantly elevated compared to those of WT mice maintained on a chow diet (Fig. 6A, 6C, 6E), consistent with the decreased neutral lipid staining (Fig 5A-D). In agreement with the enlarged liver mass and distinct color change, hepatic cholesterol was dramatically increased in the western fed wild type and V1KO mice. Consistent with the Oil Red O staining, V1KO mice showed a greater total accumulation of hepatic cholesterol than WT mice on the Western diet (Fig. 3A). This was similarly accompanied by an increase in triglycerides and hepatic free fatty acid contents for both WT and V1KO mice on the western diet (Fig. 6C-F). Increases in triglycerides and hepatic free fatty acid contents following western diet regimen were more pronounced in male V1KO mice compared to WT counterparts. Interestingly, this pronounced increase in lipid content is lost in female V1KO mice on the Western diet, indicating there may be sex-specific differences in cholesterol and lipid homeostasis.

**Fig. 6:**
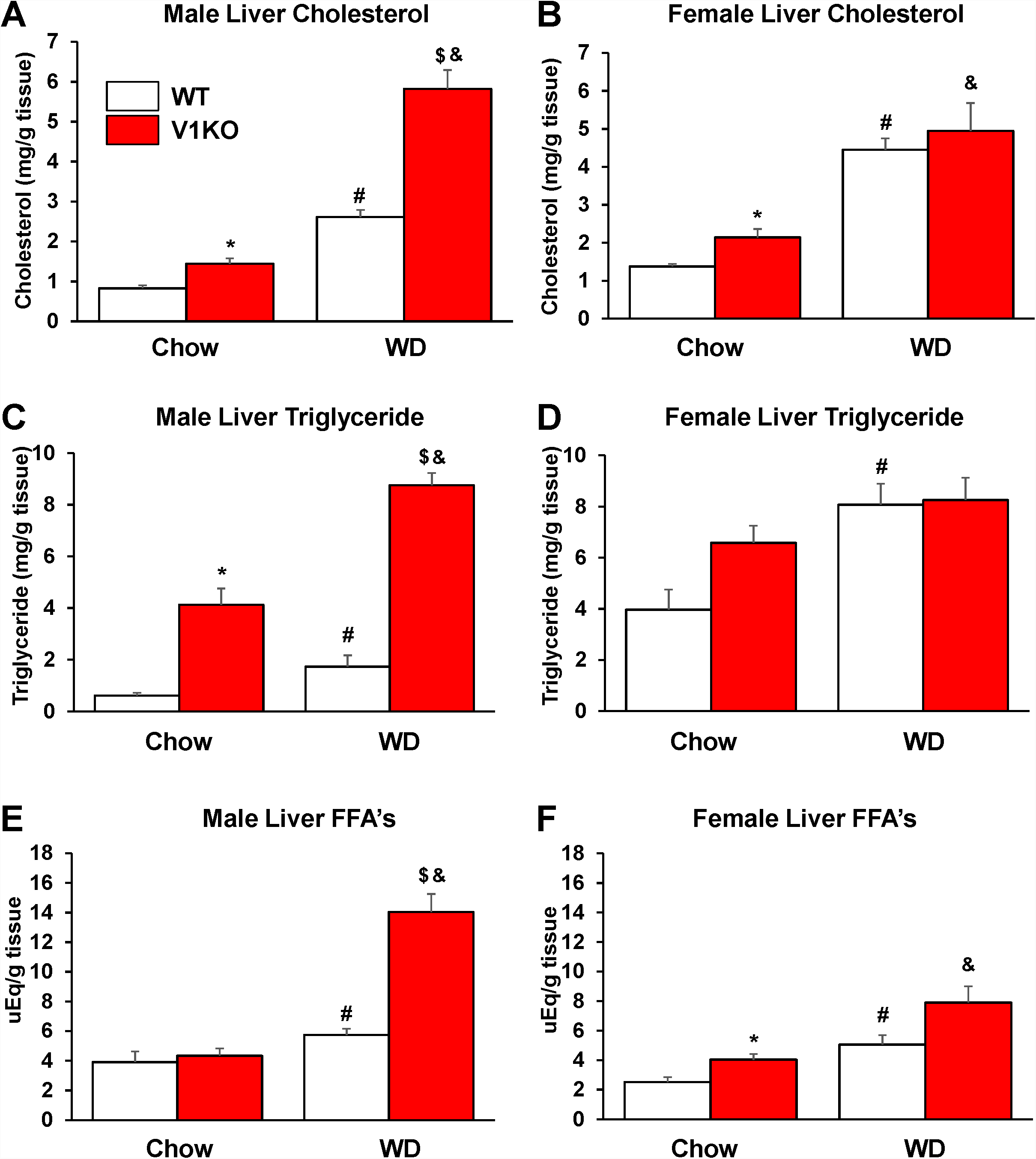
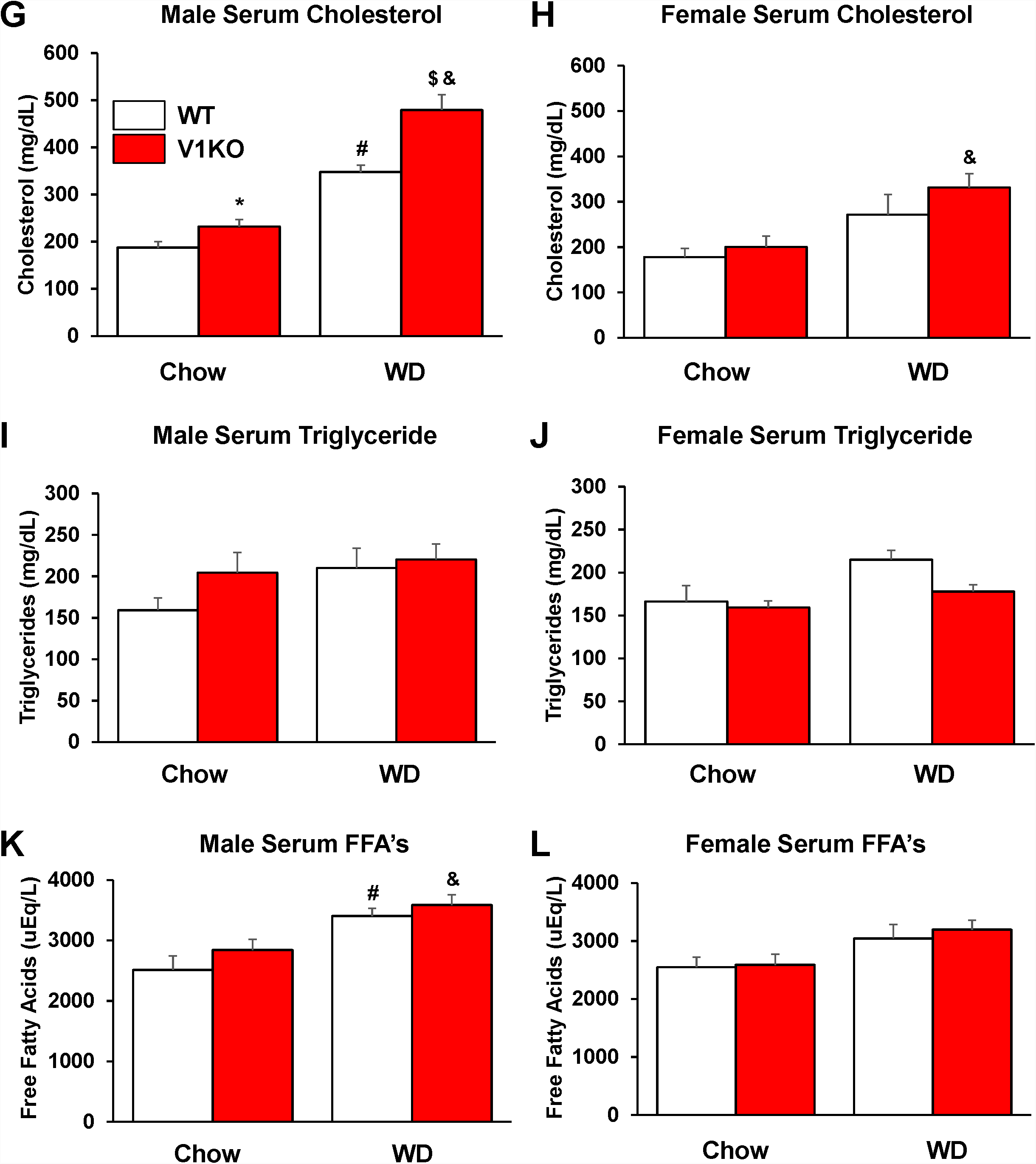
Altere lipi handling V1KO mice fe standar chow vs. western diets. Sex-specific hepatic and serum cholesterol (A/B and G/H), triglycerides (C/D and I/J), and free fatty acid (FFA; E/F and K/L) concentrations were measured in WT and V1KO mice. All values are expressed as mean ± SEM, Symbol indicates *P* < 0.05 for: *= genotype effect within chow diet. #=dietary effect with WT genotype. $=genotype effect within WD. & = dietary effect within knockout phenotype.

Interestingly, although there were dramatic and differential alterations in hepatic lipid contents in both wild type and V1KO mice following consumption of the different diets, serum cholesterol levels were markedly elevated lipid levels whereas serum TG and FFA levels were relatively unchanged (Fig. 6I).

## 4. Discussion

This study was designed to delineate the mechanisms by which TRPV1 channels regulate hepatic function and lipid metabolism. Our major findings include: 1) V1KO mice are increasingly susceptible to diet-induced obesity compared to WT mice; 2) V1KO mice exhibit severe glucose intolerance due to diet-induced obesity; 3) V1KO mice display augmented liver mass and lipid accumulation compared to WT mice; 4) Hepatotoxicity in V1KO mice correlates with enhanced levels of hepatic cholesterol, triglycerides, and free fatty acids; 5) Sex differences reveal protective effects against diet-induced obesity and its associated complications in female V1KO mice. This study provides direct evidence that lack of TRPV1 leads to hepatic dysfunction via regulation of liver and serum lipid homeostasis.

Recent evidence indicates that activation of the TRPV1 cation channel by its agonist, capsaicin, is beneficial for the management of obesity, diabetes and related diseases. As there are no approved therapeutic regimes for treatment of NAFLD, there is a pressing need to search for agents that ameliorate NAFLD phenotypes. A role for TRPV1 in hepatic function has yet to be fully determined, though recent evidence suggests an ability to reduce liver lipid deposition through upregulation of UCP2 (11) and regulate of PPARα-mediated reduction in inflammatory cytokines (10) and PPARδ-dependent autophagy (13). We initially sought to understand the effects of high fat diet (diet-induced obesity) in wild-type and V1KO mice. In the current study, a 24 week western diet regimen resulted in V1KO mice having increased weight gain over the course of the 24 week period compared to their WT counterparts (Fig. 1). This is supported by recent findings (14-16), yet contrasts seminal findings which reported that V1KO mice were resistant to diet-induced obesity (17, 18). A key difference, however, can be attributed to the fat content in the diets used in the current study compared to Motter and Ahern (42% fat vs. 11% fat respectively). This corresponding enhanced weight gain was also evident in female V1KO mice fed the western diet; yet at the end of the diet regimen, female V1KO mice weighed less than their male counterparts (40.9 ± g vs *35.5 ± g) suggesting females were afforded some degree of protection against diet-induced obesity (Fig. 1B). Notably, western fed WT female mice did not display significant alterations of body weight when compared to their chow fed counterparts over the course of the 24-week period. These findings, although paradoxical, are consistent with previous investigations (19-21) suggesting that particular strains of female mice may be protected from diet-induced obesity.

A role for TRPV1 in the regulation of adipocytes has been previously described (22-24). Additionally, our data and others (14) support the notion that the loss of TRPV1 and resulting susceptibility to diet-induced obesity leads to a further diminished “lean mass:body weight ratio” and enhanced whole body fat mass (as measured by EchoMRI) in V1KO mice fed the western diet (Fig. 1B and 1C) compared to WT counterparts.

While it is largely accepted that TRPV1 contributes to overall metabolic homeostasis via roles in thermogenesis, regulation of adipogenesis, satiety and energy expenditure (7, 25), it is increasingly apparent that the role of TRPV1 is much more complex. For instance, TRPV1 signaling has been implicated in enhancing fat accumulation to thus modulate energy and glucose homeostasis (26). In this regard, TRPV1 has been shown to influence glucagon-like peptide-1 (GLP-1) secretion (27), adipogenesis (23), and innervation of the liver to regulate glucose handling (28). Furthermore, previous investigations reported that elimination of capsaicin-sensitive fibers in Zucker diabetic fatty rats led to improved glucose tolerance and insulin resistance (29). Complementary experiments illustrated the ability of TRPV1 antagonism to reverse the elevated glucose and insulin levels and improve glucose tolerance in a diabetic mouse model (30). TRPV1 has also been linked to insulin secretion regulation and islet inflammation through pancreatic islets innervation (31). However, the uncertainty surrounding the ability of TRPV1 to regulate insulin and glucose handling needs further investigation. A large body of evidence supports a complex interaction between NAFLD and insulin resistance (32, 33). Our current findings further substantiate the role for TRPV1 in glucose handling and insulin sensitivity as our data indicates an increased fasting glucose level and glucose intolerance following the high-fat diet regimen in V1KO mice compared to high-fat diet fed WT mice. Importantly, while both of these parameters are increased in the high-fat diet fed wild-type mice, the effects are further amplified by the absence of TRPV1. Female mice fed a chow diet are not glucose intolerant, regardless of genotypic background. The differences in these observations between sexes indicate that female V1KO mice are comparable to wild-type mice on a normal diet, while male V1KO mice are pre-dispositioned to dysfunction without Western diet.

Recent research has begun to elucidate the mechanisms that govern TRPV1-dependent hepatic regulation of lipid homeostasis. For instance, a role for TRPV1-mediated increases in hepatic UCP2 expression has been demonstrated, which may account for decreased fatty deposition following a high fat diet (11). Additional investigations also demonstrated a role for attenuated NAFLD via enhanced PPARδ-mediated autophagy enhancement (13). In the current study, western diet fed WT and V1KO mice exhibited significant increases in liver weights and lipid accumulation. Furthermore, Western diet resulted in augmentation in liver size in both WT and V1KO mice, although the severity of these symptoms was more developed in the V1KO mice. We next examined the cellular indices of NAFLD using Oil Red O staining. As expected from the gross liver morphology, lipid accumulation was particularly dramatic in the V1KO Western-fed livers (Fig. 5A-D). Lipid drop accumulation as seen by increased adipocyte number and size were evident in V1KO animals. Importantly, these are key characteristics attributed to NAFLD.

Similar observations were made in both WT and V1KO animals with H&E staining; hepatocyte damage and degeneration, increased intercellular sinusoidal spaces, inflammation, increased lipid droplet size and number and disruption of overall architecture was clearly evident (Fig. 5E-H). The severity of these changes were strikingly more evident in V1KO mice fed the western diet, confirming a role for TRPV1 in diet-induced hepatotoxicity and lipid accumulation.

To correlate the increase in lipid accumulation observed using Oil Red O staining, we next examined liver lipid content in respect to whole liver cholesterol, triglyceride and free fatty acid levels. 24 weeks of Western diet resulted in higher total hepatocyte cholesterol, triglyceride and free fatty acid levels in both WT and V1KO mice, however, this effect was magnified in V1KO mice, leading to significant differences between groups. While Western diet increased whole liver cholesterol, triglyceride and free fatty acid levels in females, there was no measurable difference between genotypes. In addition, similar measurements taken from serum revealed different trends than that of the liver. The effects of genotype, sex, or diet elicited no impact on serum triglyceride and free fatty acid levels.

### 4.1. Conclusions

In conclusion, the current data demonstrates a role for TRPV1 in diet-induced obesity and subsequent induction of NAFLD. Western fed V1KO mice, when compared to their WT counterparts, demonstrate various pathophysiological disturbances including glucose intolerance and exacerbated progression of diet-induced obesity, as well as altered lipid profiles and gross morphological alterations in liver composition and size. Furthermore, there appears to be a sex-specific component whereby female V1KO mice were resistant to diet-induced obesity and the progression of NAFLD when compared to the male mice. Therefore, it is evident that the activation of TRPV1 may lead to improved hepatic health and prevention of NAFLD, yet the need to explore the role of TRPV1 in the liver and its contribution to the onset and development of NAFLD continues to grow.

## Acknowledgments

We would like to thank Dr. John Chiang (Northeast Ohio Medical University) and Dr. Maureen Gannon (Vanderbilt University Medical Center) for helpful discussions and critical reading of the manuscript. We would also like to thank members of the Chiang lab for assistance with technical training.

## Author Contributions

IB, PC, and DSD participated in research design and analysis. PC, IB, SA, LE, DJD, JNF, JMF and PP conducted the experiments. IB, PC, and SA wrote or contributed to the writing of the manuscript.

## Non-standard abbreviations

T2DM: Type 2 diabetes mellitus
NAFLD: Nonalcoholic fatty liver disease
TRPV1: Transient receptor potential channel vanilloid subtype 1
GTT: glucose tolerance test
RER: respiratory exchange ratio

